# Common brain areas for processing physiologically and socially ‘needed’ stimuli

**DOI:** 10.1101/2022.09.30.510041

**Authors:** J. Bosulu, Y. Luo, S. Hétu

## Abstract

We looked at the overlap between brain areas related to perception of physiologically and socially (non-physiological) needed stimuli and how they might regulate serotonin levels. First, we conducted separate ALE meta-analyses on published results pertaining to brain activation patterns when participants perceived food while hungry or water while thirsty, and social interactions while being excluded. This allowed us to identify common consistent brain activation patterns for physiological and social needed stimuli. We also looked at significant spatial association between the common network and serotonin receptor distribution. We found that regions within the mid-posterior insula, the anterior cingulate cortex and the caudate are at the intersection of physiological (hunger and thirst) and social (exclusion) aspects of ‘needing’. Furthermore, we found a significant positive spatial correlation between that common network and 5HT4 receptor among serotonin receptors. While this was the highest for serotonin receptors, it was not the highest of all receptors. Our study suggests there is a common brain pattern during the processing of physiologically and socially needed stimuli, and discusses their spatial association with serotonin receptors and its possible implication.

## INTRODUCTION

Needs are related to states of deprivation of a biologically significant stimuli or events (Bouton, 2016). They can be related to physiologically relevant stimuli and events such as food and eating for hunger, or water and drinking for thirst, but also to social stimuli and events like belongingness for social exclusion and meaningful social contact for social isolation (Maslow, 1943; Baumeister & Leary, 1995; Maner et al., 2007). Not responding to a need might lead to some adverse consequences such as physiological or psychological suffering which go beyond mere frustration (MacGregor 1960; Baumeister & Leary, 1995). Maslow (1943) (likely influenced by Hull’s (1943) drive theory) was convinced that there are motivations which are driven by basic needs instead of external rewards. He ranked these needs which later became Maslow’s pyramid. At the base, there are needs that are physiological, then needs of security, belonging, esteem, (more social needs) and actualization. When the lower order needs are satisfied to a certain extent, the higher ones appear gradually. Despite the fact that evidences seem to go against some definite hierarchy of needs both on an individual and on a societal level (Goebel & Brown, 1981; Hofstede, 1984; Tay & Diener, 2011), Maslow’s ideas indicate that physiological and social needs have something in common: they have some negative effect if not met, and hence motivate people to choose stimuli (or actions) that satisfy them. Indeed, in the brain, physiological needs and social needs can both alter the affective value of their relevant/significant needed stimuli (Chen et al., 2016; De Araujo, 2003). It’s important to stress that some social needs, such as social isolation, can have physiological effects almost similar to “true” physiological needs (Cacioppo et al., 2000; Tomova et al., 2020). Hence, in this study we focus on social needs that have less physiological effect, e.g. short term social exclusion. Indeed, the overlap between brain areas processing physiologically (e.g., through hunger and thirst) and socially (e.g., through short term social exclusion) needed stimuli have rarely been tested, especially on a meta-analytic level, which would provide a quantitative comparison of the processing of physiologically vs. socially needed stimuli.

The idea of needs can be considered to include two components : the state itself and the (needed) stimulus that alleviate that state. Since processing of the needed stimuli are more likely to control behavior, either as cues or outcomes, than the deprivation states themselves (see Bindra, 1974; Toates, 1994), we will focus on brain responses associated with this processing. Hence, here we conceptualize the brain response to the perception of a stimulus, when deprived of it, as ‘needing’. For hunger and thirst, brain studies that measured brain response to (viewing, tasting, or smelling) food or (either ingesting drops of water or viewing beverages) while hungry or thirsty have reported activations in the insula (for hunger/food: van der Lan et al., 2011; Goldstone, et. al., 2009; Siep et. al., 2009; for thirst/water: De Araujo et al., 2003; Egan et al., 2003; Becker et al., 2017; Farrell et al., 2011), orbitofrontal cortex (OFC) (for hunger/food: van der Lan et al., 2011; Goldstone, et. al., 2009; Siep et. al., 2009; Führer et. al., 2008; for thirst/water: De Araujo et al., 2003; Saker et al., 2014), dorsal striatum (for hunger/food: van der Laan, et. al., 2011; Siep et. al., 2009), anterior cingugale cortext (ACC) (for hunger/food: Goldstone, et. al., 2009; Siep et. al., 2009; Führer et. al., 2008; for thirst/water: De Araujo et al., 2003; Becker et al., 2015, Becker et al., 2017; Farrell et al., 2011; Saker et al., 2013), amygdala and parahippocampal gyrus (for hunger/food: van der Laan, et. al., 2011; LaBar et. al., 2001; Führer, 2008; Goldstone, 2009; Mohanty, et. al., 2008; Chen et. al., 2020; for thirst/water: Becker, 2015), and posterior cingulate cortex (PCC) (for thirst/water: Farrell et al., 2011). Hence, although the brain networks supporting different types of physiological needs (here hunger and thirst) are not the same, they partially overlap in the ACC and amygdala, as well as OFC and the insular cortex; specifically the mid-posterior insula which has been shown to code for needed stimuli (see Bosulu et al., 2022; Livneh et al., 2020; Livneh et al., 2017). Furthermore, the insular cortex and ACC are viewed as common regions for conscious perception of both hunger and thirst (Mckinley et al., 2019), suggesting that they are core regions for homeostatic related perception and motivation (Craig, 2003).

Studies and meta-analyses of social exclusion/belongingness that have looked at brain responses to viewing others interact, or viewing (pleasant and/or close other) social stimuli while excluded from social interaction have reported activity in the ACC (Eisenberger, et al., 2003; Masten et al., 2009; Bolling et al., 2011; Vijayakumar et al., 2017; Mwilambwe-Tshilobo & Spreng, 2021), the posterior cingulate cortex (PCC) (Vijayakumar et al., 2017; Mwilambwe & Spreng, 2021); the insula (Masten et al., 2009; Bolling et al., 2011; Mwilambwe & Spreng, 2021), the ventral striatum (Masten et al., 2009; Vijayakumar et al., 2017), and the OFC (Cacioppo et al., 2013; Vijayakumar et al., 2017). Overall, looking at these various results, social exclusion related responses to social cues/interactions seem to activate the insula, ACC, and OFC.

Overall, these different results suggest that brain networks supporting physiological and non physiological social needing at least partially overlap. Indeed, qualitatively the activation patterns of hunger/food, thirst/water, and social exclusion/belongingness often show activity within the insula (e.g. van der Lan et al., 2011; De Araujo et al., 2003; Mwilambwe-Tshilobo & Spreng, 2021; Tomova et al., 2020), ACC (ex. Goldstone, et. al., 2009; De Araujo et al., 2003; Mwilambwe-Tshilobo & Spreng, 2021) and OFC (ex. van der Lan et al., 2011; De Araujo et al., 2003; Vijayakumar et al., 2017). However, in a recent study directly testing for common brain areas when participants observed food versus social interactions, after respectively 10 hours of hunger or 10 hours of social isolation, only activations of regions containing mesolimbic and nigrostriatal dopamine neurons of the ventral tegmental area (VTA) and substantia nigra (SN) were found (Tomova et al., 2020). No study has compared physiological needs and social needs that have less physiological effect, such as short term social exclusion. Hence there is still some uncertainty with regard to if and which regions may be recruited by both the processing of physiological (hunger and thirst) and social needs (exclusion) that have less physiological effect. Hence, it seems timely to tackle this question using a meta-analysis which provides a quantitative analysis of the existing literature.

Tomova et al. (2020) study on hunger and social isolation seems to suggest that, because VTA/SN are related to dopamine, this neurotransmitter would be a neurotransmitter commonly involved in perception of physiologically and socially needed stimuli, likely turning deprivation into reward seeking for both types of needs. Beyond dopamine, we can wonder what other neurotransmitters might be related to the processing of physiological and social needed stimuli, especially for social needs that are not related to long term physiological process such as short term social exclusion. Serotonin is a good candidate for overall need processing, as it is central for biologically important sensory events (Sizemore, 2020). Indeed, low serotonin levels in the brain have been related to higher sensitivity to food (van Galen et al., 2021) and to higher reactivity to social exclusion (Preller et al., 2015). As low serotonin is often associated with aversive processing (Dayan and Huys, 2009) (which characterize deprivation); serotonin secretion is said to indicate how beneficial the current state is (Luo et al., 2016; Liu et al., 2020). In that sense, (low) serotonin level might signal states of deprivation of both biologically and socially significant stimuli/events. However such a mechanism has not been elucidated.

Hence, the present study aimed at investigating the common and specific brain activation patterns for, on one hand, the processing of physiologically needed stimuli and on the other socially needed stimuli. As an exploratory objective, to help us understand how the brain processes relevant stimuli in a deprived state, we also tested for the spatial correlation between areas recruited during both the processing for physiologically and socially needed stimuli and brain regions associated with the spatial distribution of neurotransmitter receptors, with a specific interest in serotonin receptors, given its inherent link with well-being (see Luo et al., 2016; Liu et al., 2020).

## METHODOLOGY

### Meta-analysis of brain coordinate

We used a meta-analytic approach to quantitatively assess brain activation patterns of both physiological and social aspects of ‘needing’ using large collections of data. Meta-analyses make it possible to investigate questions across different paradigms, samples and analysis approaches. Specifically, we first conducted two meta-analyses to quantitatively summarize results from functional magnetic resonance imaging (fMRI) published studies on physiological needs: hunger/food and thirst/water (Physiological-Need); when participants perceived food while hungry or water while thirsty (that we will refer to as Physiological-Need); and on social non physiological need: short term exclusion/interaction; when participants perceived social interactions (that s/he was supposed to be part of) while being excluded from that social interaction (that we will refer to as Social-Need). These meta-analyses will help us look at brain regions that are consistently recruited for processing of either physiologically or socially needed stimuli. Second, we used the single meta-analytic results for contrasts analysis as well as a conjunction analysis to identify differences and overlaps in consistent brain activation for physiological and social ‘needing’.

We used the PRISMA framework and the following keywords to identify articles related to hunger: (“hunger” OR “food deprivation”) AND (“fMRI”). For thirst, we used: (“thirst” OR “water deprivation”) AND (“fMRI”). For social exclusion, we used: (“social” AND “exclusion” AND “fmri”). Keywords were entered on PubMed (February 2021) for physiological needs and social needs. Additional articles were found by checking the articles references lists and review articles. For both physiological and social aspect of ‘needing’ the following inclusion criteria were used: healthy subjects; whole-brain analyses (with or without SVC), MNI or Talairach Coordinates (all Talairach coordinates were converted to MNI SPM152 in Ginger ALE using Lacanster transform; Lancaster et al., 2007); maps were corrected (or cluster level corrected); activation contrast only (see table 1 for selection criteria). The database returned 150 articles for hunger and 19 for thirst; 7 and 3 additional articles were found through other articles and reviews for hunger and thirst, respectively. In line with our objective, specific for physiological needs articles, we used two additional main criteria : 1) the participants were in a food or water deprivation state; 2) the participant was perceiving (visual, taste, odor, etc.) some food (while hungry) or water (while thirsty) stimulus. We found a total of 16 articles for hunger and 4 for thirst that met all inclusion criteria (see table 2 and 3).

**Table 1.** Selection Criteria Table indicating selection criteria. The red coloured “yes” or “no” means the criterion is crucial for the definition of either ‘physiological or social needing.

| Criteria | Hunger and Thirst | Social exclusion |
| --- | --- | --- |
| Privation contrast | Yes | Yes |
| Presence of the relevant rewarding cue | Yes | Yes |
| Passive viewing | Yes | Yes |
| Long term (in hours) deprivation implicating physiological factors | Yes | No |
| Healthy individuals only | Yes | Yes |
| MNI or Talairach Coordinates | Yes | Yes |
| Whole brain contrast (with or without SVC) | Yes | Yes |
| Corrected | Yes | Yes |
| Activation contrast only | Yes | Yes |
| Excluded: MRI and resting states; cognitive conjunction analysis; and functional connectivity results | Yes | Yes |

**Table 2.**
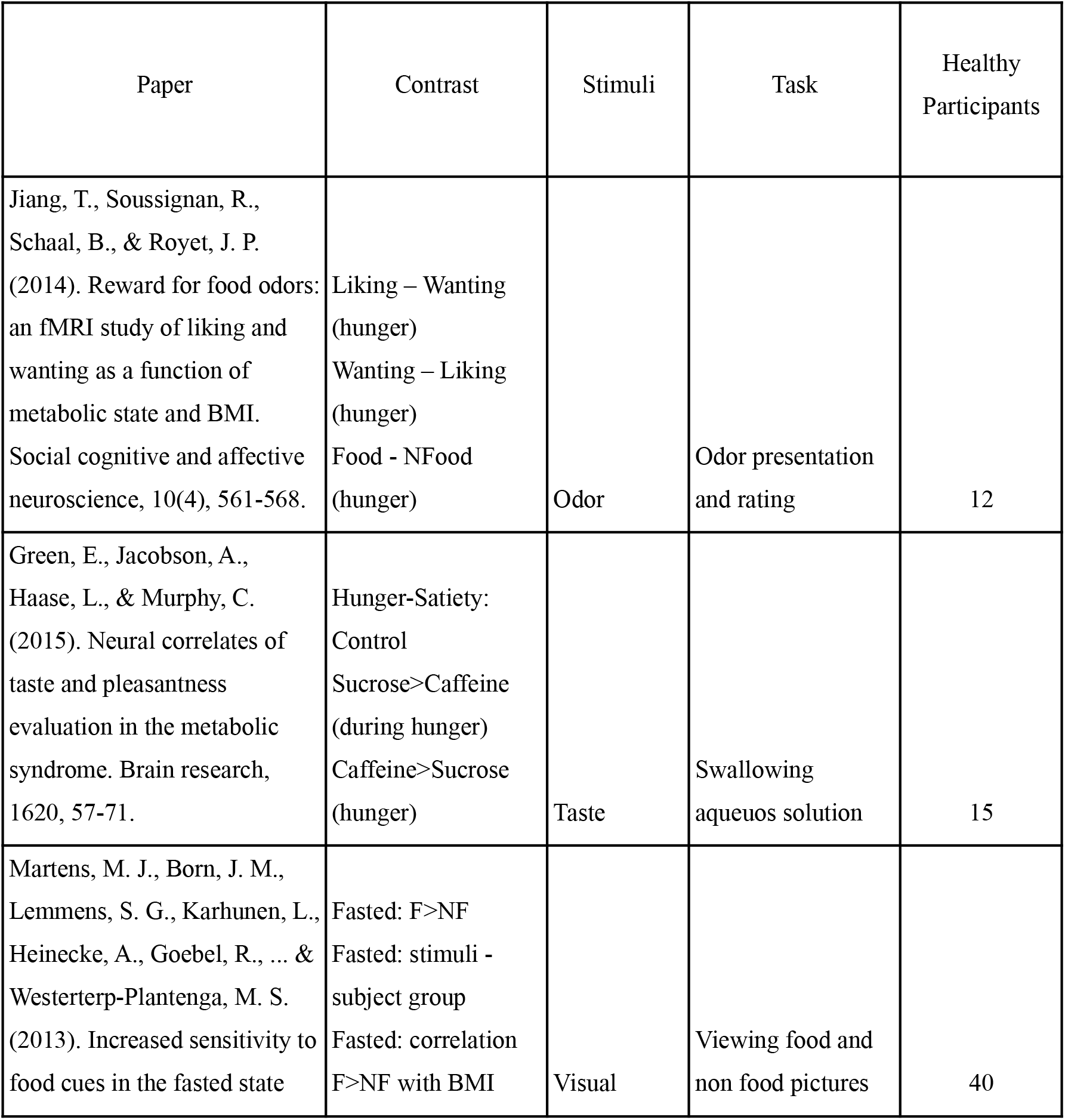
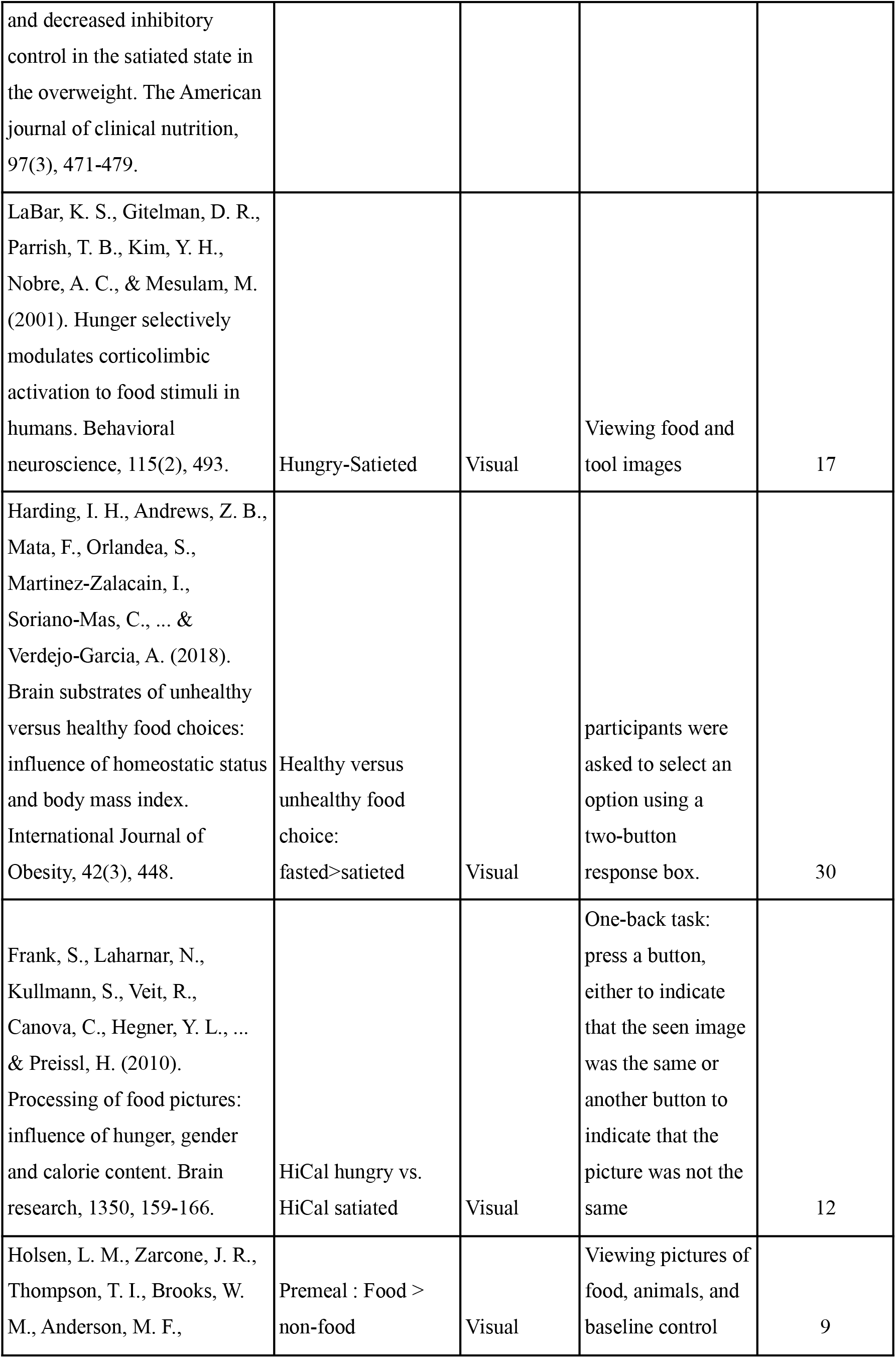
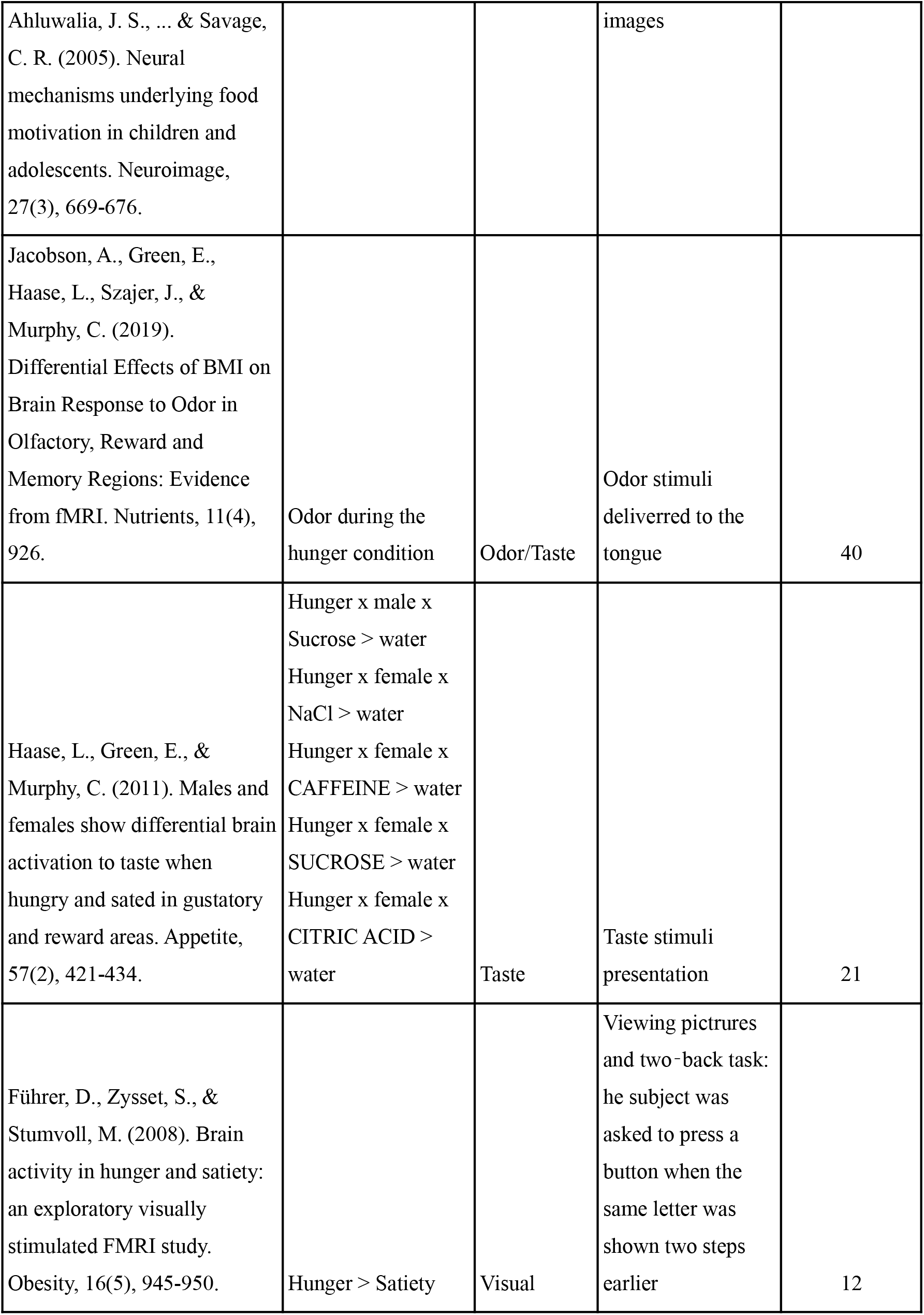
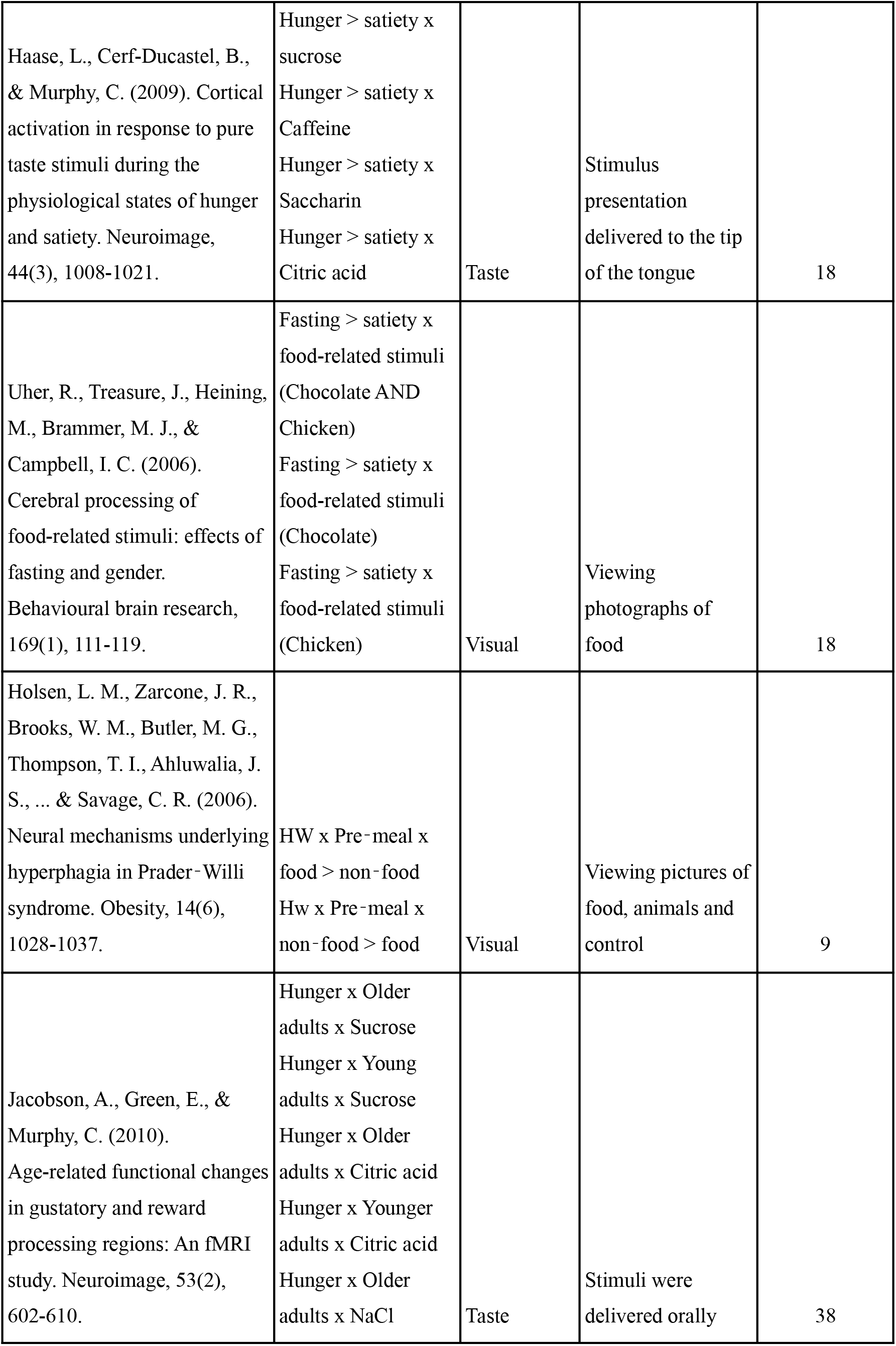
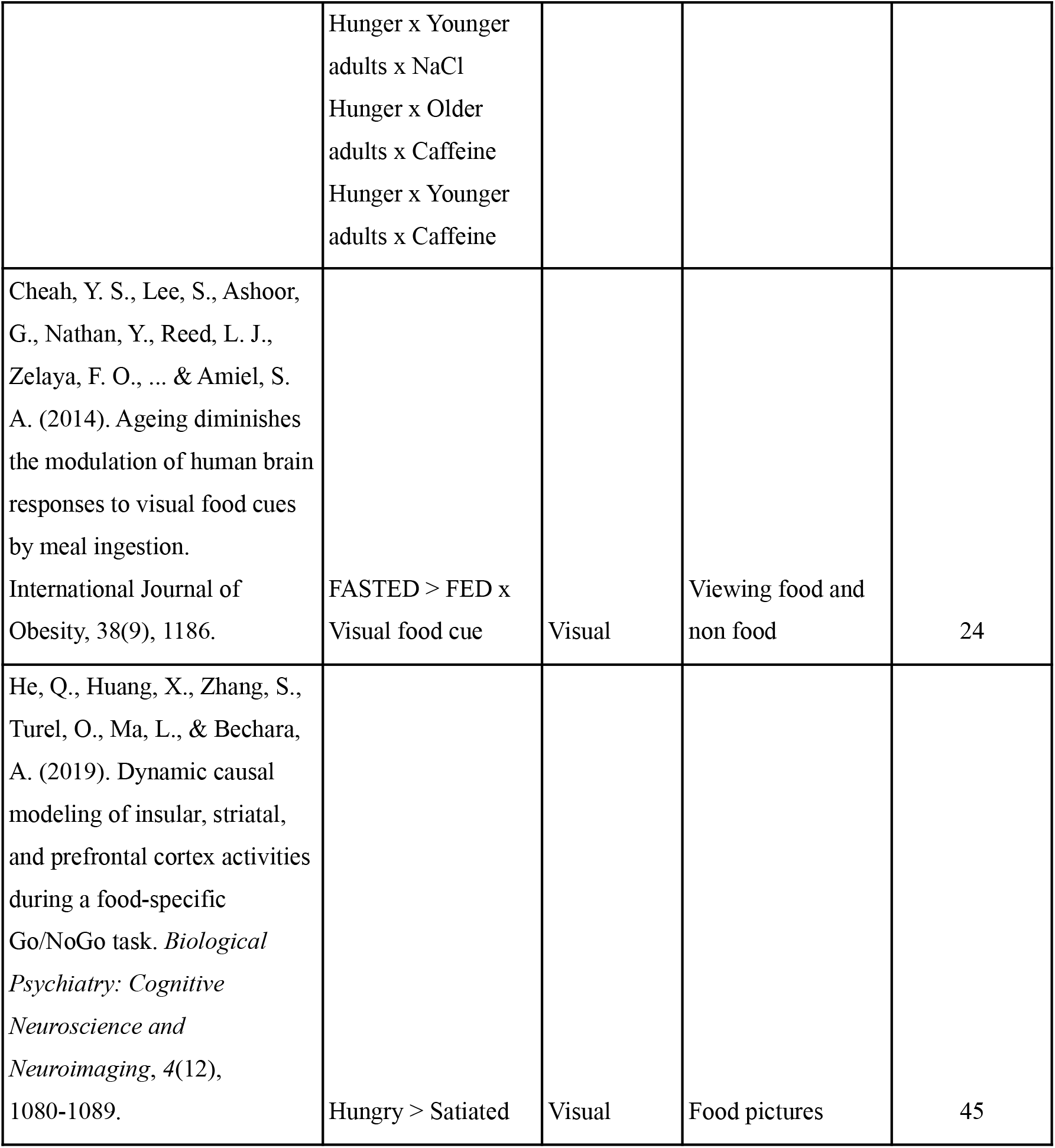
Hunger selected articles List of articles that were selected. for hunger..

**Table 3.** thirst selected articles List of articles that were selected. for thirst.

| Paper | Stimulus contrasts | Healthy participants |
| --- | --- | --- |
| Becker, C. A., Schmälzle, R., Flaisch, T., Renner, B., & Schupp, H. T. (2015). Thirst and the state-dependent representation of incentive stimulus value in human motive circuitry. *Social cognitive and affective neuroscience*, 10(12), 1722-1729. | (Beverages : Thirst > Chairs : Thirst ) vs (Beverages : No-thirst > Chairs : No-thirst ) | 24 |
| De Araujo, I. E., Kringelbach, M. L., Rolls, E. T., & McGlone, F. (2003). Human cortical responses to water in the mouth, and the effects of thirst. *Journal of neurophysiology*, 90(3), 1865-1876. | water-control water (pre and post satiety) | 11 |
| Becker, C. A., Flaisch, T., Renner, B., & Schupp, H. T. (2017). From thirst to satiety: the anterior mid-cingulate cortex and right posterior insula indicate dynamic changes in incentive value. *Frontiers in human neuroscience*, 11, 234. | (BOLD) activity proportional to the amount of water ingested | 24 |
| Saker, P., Farrell, M. J., Egan, G. F., McKinley, M. J., & Denton, D. A. (2018). Influence of anterior midcingulate cortex on drinking behavior during thirst and following satiation. *Proceedings of the National Academy of Sciences*, 115(4), 786-791. | Thirst > No-Thirst | 20 |

Regarding social exclusion/isolation, we found 129 articles, 22 were selected, and 4 more articles were found through other articles and reviews. Similarly to Physiological-Need, two additional main criteria were used for social needs : 1) the participant was in a social deprivationstate (i.e., s/he was either isolated from others or experienced social exclusion); 2) the participant was perceiving some social interaction s/he was excluded from. It’s important to note that these criteria resulted in all of our included articles for social needs using the cyber ball task (a virtual ball tossing game with other individuals from which the participant is excluded (Williams et al., 2000)), and were thus related to short term social exclusion. Social exclusion (as threat to fundamental social needs in humans) will cause emotional distress (Williams, 2007), and it has been proposed that the cyberball exclusion paradigm can induce need-like emotional distress (Bernstein and Claypool (2012). Based on these criteria and theories, we selected a total of 26 articles (for social exclusion) for the meta-analysis (see table 4).

**Table 4.** social exclusion selected articles List of articles that were selected. for social exclusion.

|  |  |  |  |
| --- | --- | --- | --- |
| Bach, P., Frischknecht, U., Bungert, M., Karl, D., Vollmert, C., Vollstädt-Klein, S., ... & Hermann, D. (2019). Effects of social exclusion and physical pain in chronic opioid maintenance treatment: fMRI correlates. *European Neuropsychopharmacology*, 29(2), 291-305. | Exclusion > Inclusion | Cyberball task | 21 |
| Bolling, D. Z., Pelphrey, K. A., & Vander Wyk, B. C. (2015). Trait-level temporal lobe hypoactivation to social exclusion in unaffected siblings of children and adolescents with autism spectrum disorders. *Developmental cognitive neuroscience*, 13, 75-83. | Social Exclusion > Fair Play | Cyberball task | 15 |
| Bolling, D. Z., Pitskel, N. B., Deen, B., Crowley, M. J., Mayes, L. C., & Pelphrey, K. A. (2011). Development of neural systems for processing social exclusion from childhood to adolescence. *Developmental science*, 14(6), 1431-1444. | comparison of social exclusion and fair play | Cyberball task | 26 |
| Bolling, D. Z., Pitskel, N. B., Deen, B., Crowley, M. J., McPartland, J. C., Kaiser, M. D., ... & Pelphrey, K. A. (2011). Enhanced neural responses to rule violation in children with autism: a comparison to social exclusion. *Developmental Cognitive Neuroscience*, 1(3), 280-294. | Social exclusion > fair play | Cyberball task | 24 |
| Bolling, D. Z., Pitskel, N. B., Deen, B., Crowley, M. J., McPartland, J. C., Mayes, L. C., & Pelphrey, K. A. (2011). Dissociable brain mechanisms for processing social exclusion and rule violation. *NeuroImage*, 54(3), 2462-2471. | Social exclusion > Fair play | Cyberball | 26 |
| Cheng, T. W., Vijayakumar, N., Flournoy, J. C., de Macks, Z. O., Peake, S. J., Flannery, J. E., ... & Pfeifer, J. H. (2020). Feeling left out or just surprised? Neural correlates of social exclusion and overinclusion in adolescence. *Cognitive, Affective, & Behavioral Neuroscience*, 1-16. | Exclusion>Inclusion | Cyberball task | 97 |
| Cogoni, C., Carnaghi, A., & Silani, G. (2018). Reduced empathic responses for sexually objectified women: An fMRI investigation. *Cortex*, 99, 258-272. | Self (exclusion > inclusion). | social pain task | 36 |
| de Water, E., Mies, G. W., Ma, I., Mennes, M., Cillessen, A. H., & Scheres, A. (2017). Neural responses to social exclusion in adolescents: Effects of peer status. *Cortex*, 92, 32-43. | Exclusion (popular + average players) > Inclusion Ball (popular + average players)<br>Exclusion (popular + average players) > Inclusion No Ball (popular + average players) | Cyberball task | 52 |
| DeWall, C. N., Masten, C. L., Powell, C., Combs, D., Schurtz, D. R., & Eisenberger, N. I. (2012). Do neural responses to rejection depend on attachment style? An fMRI study. *Social cognitive and affective neuroscience*, 7(2), 184-192. | exclusion vs inclusion | Cyberball task | 25 |
| Eisenberger, N. I., Lieberman, M. D., & Williams, K. D. (2003). Does rejection hurt? An fMRI study of social exclusion. *Science*, 302(5643), 290-292. | explicit social exclusion<br>implicit social exclusion |  | 13 |
| Falk, E. B., Cascio, C. N., O'Donnell, M. B., Carp, J., Tinney Jr, F. J., Bingham, C. R., ... & Simons-Morton, B. G. (2014). Neural responses to exclusion predict susceptibility to social influence. *Journal of Adolescent Health*, 54(5), S22-S31. | Exclusion>inclusion | Cyberball task | 36 |
| Gradin, V. B., Waiter, G., Kumar, P., Stickle, C., Milders, M., Matthews, K., ... & Steele, J. D. (2012). Abnormal neural responses to social exclusion in schizophrenia. *PloS one*, 7(8), e42608. | Activations with increasing social exclusion | Cyberball task | 20 |
| Gilman, J. M., Curran, M. T., Calderon, V., Schuster, R. M., & Evins, A. E. (2016). Altered neural processing to social exclusion in young adult marijuana users. *Biological Psychiatry: Cognitive Neuroscience and Neuroimaging*, 1(2), 152-159. | Exclusion > Fair Play | Cyberball task | 42 |
| Gonzalez, M. Z., Beckes, L., Chango, J., Allen, J. P., & Coan, J. A. (2015). Adolescent neighborhood quality predicts adult dACC response to social exclusion. *Social cognitive and affective neuroscience*, 10(7), 921-928. | Exclusion > inclusion | Cyberball task | 85 |
| Le, T. M., Zhornitsky, S., Wang, W., & Li, C. S. R. (2020). Perceived burdensomeness and neural responses to ostracism in the Cyberball task. *Journal of Psychiatric Research*, 130, 1-8. | Activations to social exclusion. | Cyberball task | 64 |
| Luo, S., Yu, D., & Han, S. (2016). Genetic and neural correlates of romantic relationship satisfaction. *Social cognitive and affective neuroscience*, 11(2), 337-348. | social exclusion vs inclusion | Cyberball task | 42 |
| Moor, B. G., Güroğlu, B., de Macks, Z. A. O., Rombouts, S. A., Van der Molen, M. W., & Crone, E. A. (2012). Social exclusion and punishment of excluders: neural correlates and developmental trajectories. *Neuroimage*, 59(1), 708-717. | No Ball- exclusion game<br>> Ball- inclusion game<br>No Ball- exclusion game<br>> No Ball- inclusion game | Cyberball game | 51 |
| Nishiyama, Y., Okamoto, Y., Kunisato, Y., Okada, G., Yoshimura, S., Kanai, Y., ... & Yamawaki, S. (2015). fMRI Study of social anxiety during social ostracism with and without emotional support. *PloS one*, 10(5), e0127426. | Exclusion -Inclusion | Cyberball task | 46 |
| Novembre, G., Zanon, M., & Silani, G. (2015). Empathy for social exclusion involves the sensory-discriminative component of pain: a within-subject fMRI study. *Social cognitive and affective neuroscience*, 10(2), 153-164. | Main effect of social pain: Self (Exclusion > Inclusion) | Cyberball task | 23 |
| Puetz, V. B., Kohn, N., Dahmen, B., Zvyagintsev, M., Schüppen, A., Schultz, R. T., ... & Konrad, K. (2014). Neural response to social rejection in children with early separation experiences. *Journal of the American Academy of Child & Adolescent Psychiatry*, 53(12), 1328-1337. | Controls SoEx>TechEx | Cyberball task | 26 |
| Radke, S., Seidel, E. M., Boubela, R. N., Thaler, H., Metzler, H., Kryspin-Exner, I., ... & Derntl, B. (2018). Immediate and delayed neuroendocrine responses to social exclusion in males and females. *Psychoneuroendocrinology*, 93, 56-64. | Social exclusion > Inclusion | Social exclusion task and cover story (ball tossing game) | 80 |
| van der Meulen, M., Steinbeis, N., Achterberg, M., van IJzendoorn, M. H., & Crone, E. A. (2018). Heritability of neural reactions to social exclusion and prosocial compensation in middle childhood. *Developmental cognitive neuroscience*, 34, 42-52. | Exclusion > Inclusion | Prosocial Cyberball Game | 283 |
| Wagels, L., Bergs, R., Clemens, B., Bauchmüller, M., Gur, R. C., Schneider, F., ... & Kohn, N. (2017). Contextual exclusion processing: an fMRI study of rejection in a performance-related context. *Brain imaging and behavior*, 11(3), 874-886. | Exclusion > Inclusion | mental visualisation abilities while playing a ball-tossing game | 40 |
| Will, G. J., van Lier, P. A., Crone, E. A., & Güroğlu, B. (2016). Chronic childhood peer rejection is associated with heightened neural responses to social exclusion during adolescence. *Journal of abnormal child psychology*, 44(1), 43-55. | Social exclusion (Exclusion: not receiving the ball > Inclusion: receiving the ball) Social exclusion (Exclusion: not receiving the ball > Inclusion: not receiving the ball) | Cyberball task | 44 |
| Chester, D. S., Lynam, D. R., Milich, R., & DeWall, C. N. (2018). Neural mechanisms of the rejection–aggression link. *Social cognitive and affective neuroscience*, 13(5), 501-512. | Rejection > Acceptance | Cyberball task | 60 |
| Wudarczyk, O. A., Kohn, N., Bergs, R., Gur, R. E., Turetsky, B., Schneider, F., & Habel, U. (2015). Chemosensory anxiety cues moderate the experience of social exclusion—an fMRI investigation with Cyberball. *Frontiers in psychology*, 6, 1475. | Exclusion > Inclusion<br>for: (a) sports<br>chemosensory condition; | Cyberball task | 24 |

Meta-analyses were conducted with the activation likelihood estimation (ALE) approach using the Brainmap’s GingerALE application. The revised ALE meta-analysis by Eickhoff and colleagues (2009) treats activation foci not as single point, but as spatial probability distributions centered at the given coordinates (Eickhoff et. al, 2012). It models spatial uncertainty by using an estimation of the inter-subject and inter-laboratory variability (typically observed in neuroimaging experiments). An ALE map is obtained by computation of union of activation probabilities for each voxel of all included experiments; and a permutation procedure is used to test for true convergence vs. random clustering (Eickhoff, et. al, 2012). The inference is done through the use of random-effects analysis that calculates the above-chance clustering between experiments. Furthermore, the algorithm gives more weight to gray matter compared to white matter by limiting the meta-analysis to an anatomically constrained space specified by a gray matter mask. For each single meta-analysis, we used the MNI152 coordinate system and the less conservative (larger) mask size. For hunger and thirst, there were 20 articles, 44 experiments, 856 subjects and 612 foci. (Hunger and thirst were merged together as physiological ‘needing’). For social exclusion, there were 26 articles, 33 experiments, 1511 subjects and 342 foci. In our study, for main individual meta-analyses, all maps were thresholded using a cluster-level family-wise error (cFWE) correction (P < 0.05) with a cluster-forming threshold of P < 0.001(uncorrected at the voxel level) (Eklund et al., 2016; Woo et al., 2014), and 1000 permutations. The contrasts analyses ([Physiological-Need] > [Social-Need] and [Social-Need] > [Physiological-Need]) compared the two different datasets (i.e. the ALE results from the Physiological-Need and Social-Need meta-analyses) for statistically significant differences. The conjunction analysis ([Physiological-Need] AND [Social-Need]), which is the main purpose of this study, was performed by intersecting the thresholded maps for physiological and social needs and allowed us to identify potential brain areas that were consistently activated during both physiological and social needs. Maps from meta-analyses were overlaid on a MNI template using Mango (http://ric.uthscsa.edu/mango/).

### Spatial correlation with neurotransmitters

Using the conjunction results ([Physiological-Need] AND [Social-Need]), we also looked at possible spatial (topographical) relationships between this intersection map and neurotransmitters distribution whole-brain maps available in the JuSpace (as of december 2021). The latter is a MATLAB based toolbox introduced by Dukart and colleagues (2021) which allows for cross-modal correlation of spatial patterns of MRI or fMRI based measures with PET derived biological distribution of specific tissue properties, i.e. receptor density estimates covering dopaminergic, serotonergic, noradrenergic receptors and/or transporters (Dukart et al., 2021). For the purpose of this study, we focused on serotonin receptors. Significant spatial association between subregions of the conjunction results and each neurotransmitter map were examined by comparing the distribution of z-transformed correlations (adjusted for spatial autocorrelation (i.e. adjusting for local gray matter probabilities as estimated from TPM.nii provided with SPM12)) against null distribution using one-sample t-tests (Dukart et al., 2021).

## RESULTS

### Single meta-analyses

The single meta-analysis on Physiological-Need, i.e. food and water perception during hunger & thirst (combined), revealed consistent activation within the following regions: the bilateral anterior insula, right middle insula, bilateral posterior insula, right claustrum, right putamen, bilateral ACC (B24 and B25), the bilateral caudate head, right caudate body, the left parahippocampal gyrus, the left medial frontal gyrus, right amygdala, right uncus, right mamillary body and right hippocampus (see table 5 and figure 1).

**Table 5.** Hunger & thirst Coordinates for peak activated clusters in the hunger & thirst condition. We used a cluster-level family-wise error (cFWE) correction (P < 0.05) with a cluster-forming threshold of P < 0.001(uncorrected at the voxel level) (Eklund et al., 2016; Woo et al., 2014), and 1000 permutations. cluster-level family-wise error (cFWE) correction (P < 0.05) with a cluster-forming threshold of P < 0.001(uncorrected at the voxel level) (Eklund et al., 2016; Woo et al., 2014), and 1000 permutations.

| Cluster # | x | y | z | ALE | P | Z | Label (Nearest Gray Matter within 5mm) |
| --- | --- | --- | --- | --- | --- | --- | --- |
| 1 | 38 | 10 | 6 | 0.031111395 | 2.5334245E-8 | 5.448993 | Right Cerebrum.Sub-lobar.Insula.Gray Matter.Brodmann area 13 |
| 1 | 42 | 4 | -10 | 0.02957781 | 7.35114E-8 | 5.2562838 | Right Cerebrum.Sub-lobar.Insula.Gray Matter.Brodmann area 13 |
| 1 | 40 | -4 | 2 | 0.027335959 | 3.4063015E-7 | 4.966653 | Right Cerebrum.Sub-lobar.Clastrum.Gray Matter.* |
| 1 | 46 | -12 | 4 | 0.02393658 | 3.1988484E-6 | 4.51281 | Right Cerebrum.Sub-lobar.Insula.Gray Matter.Brodmann area 13 |
| 1 | 34 | -12 | 6 | 0.02189089 | 1.19065735E-5 | 4.2257686 | Right Cerebrum.Sub-lobar.Lentiform Nucleus.Gray Matter.Putamen |
| 1 | 34 | 20 | -6 | 0.021366583 | 1.654994E-5 | 4.1510224 | Right Cerebrum.Sub-lobar.Clastrum.Gray Matter.* |
| 1 | 28 | -8 | 0 | 0.019493565 | 5.3015683E-5 | 3.8763602 | Right Cerebrum.Sub-lobar.Lentiform Nucleus.Gray Matter.Putamen |
| 2 | -38 | 0 | -2 | 0.031286273 | 2.233257E-8 | 5.4713836 | Left Cerebrum.Sub-lobar.Clastrum.Gray Matter.* |
| 2 | -42 | -12 | 10 | 0.022954691 | 6.046834E-6 | 4.375899 | Left Cerebrum.Sub-lobar.Insula.Gray Matter.Brodmann area 13 |
| 2 | -40 | 8 | 12 | 0.017407449 | 1.8151532E-4 | 3.565596 | Left Cerebrum.Sub-lobar.Insula.Gray Matter.Brodmann area 13 |
| 3 | 14 | 36 | -4 | 0.022025552 | 1.0891327E-5 | 4.245793 | Right Cerebrum.Limbic Lobe.Anterior Cingulate.Gray Matter.Brodmann area 24 |
| 3 | 10 | 24 | 0 | 0.021441728 | 1.5834556E-5 | 4.1611233 | Right Cerebrum.Sub-lobar.Caudate.Gray Matter.Caudate Head |
| 3 | 8 | 20 | -12 | 0.02108268 | 1.985968E-5 | 4.1091094 | Right Cerebrum.Limbic Lobe.Anterior Cingulate.Gray Matter.Brodmann area 25 |
| 3 | 14 | 10 | 6 | 0.018935341 | 7.378362E-5 | 3.7951293 | Right Cerebrum.Sub-lobar.Caudate.Gray Matter.Caudate Body |
| 3 | 10 | 14 | -2 | 0.0177747 | 1.4716572E-4 | 3.620239 | Right Cerebrum.Sub-lobar.Caudate.Gray Matter.Caudate Head |
| 4 | -26 | 2 | -18 | 0.024686959 | 1.9669078E-6 | 4.61486 | Left Cerebrum.Limbic Lobe.Parahippocampal Gyrus.Gray Matter.Brodmann area 34 |
| 4 | -14 | 18 | -20 | 0.020973139 | 2.128351E-5 | 4.093089 | Left Cerebrum.Frontal Lobe.Medial Frontal Gyrus.Gray Matter.Brodmann area 25 |
| 4 | -14 | 22 | -12 | 0.017613713 | 1.6152841E-4 | 3.5960736 | Left Cerebrum.Sub-lobar.Caudate.Gray Matter.Caudate Head |
| 4 | -6 | 22 | -10 | 0.015441236 | 5.697342E-4 | 3.2536144 | Left Cerebrum.Limbic Lobe.Anterior Cingulate.Gray Matter.Brodmann area 24 |
| 5 | 26 | 6 | -16 | 0.02176692 | 1.2846617E-5 | 4.2086267 | Right Cerebrum.Sub-lobar.Lentiform Nucleus.Gray Matter.Putamen |
| 5 | 30 | -10 | -20 | 0.02068114 | 2.5509367E-5 | 4.0509133 | Right Cerebrum.Limbic Lobe.Parahippocampal Gyrus.Gray Matter.Amygdala |
| 5 | 32 | -4 | -26 | 0.019510057 | 5.2373212E-5 | 3.8793273 | Right Cerebrum.Limbic Lobe.Parahippocampal Gyrus.Gray Matter.Amygdala |
| 5 | 34 | 4 | -28 | 0.019396378 | 5.597474E-5 | 3.8631182 | Right Cerebrum.Limbic Lobe.Uncus.Gray Matter.Brodmann area 28 |
| 6 | 14 | -16 | 0 | 0.027833832 | 2.4450569E-7 | 5.030599 | Right Cerebrum.Sub-lobar.Thalamus.Gray Matter.Mammillary Body |
| 7 | 32 | -38 | -4 | 0.021315122 | 1.7078451E-5 | 4.1438227 | Right Cerebrum.Temporal Lobe.Sub-Gyrus.Gray Matter.Hippocampus |
| 7 | 36 | -22 | -4 | 0.02061068 | 2.6645188E-5 | 4.0407095 | Right Cerebrum.Sub-lobar.Lentiform Nucleus.Gray Matter.Putamen |

**Figure 1.**
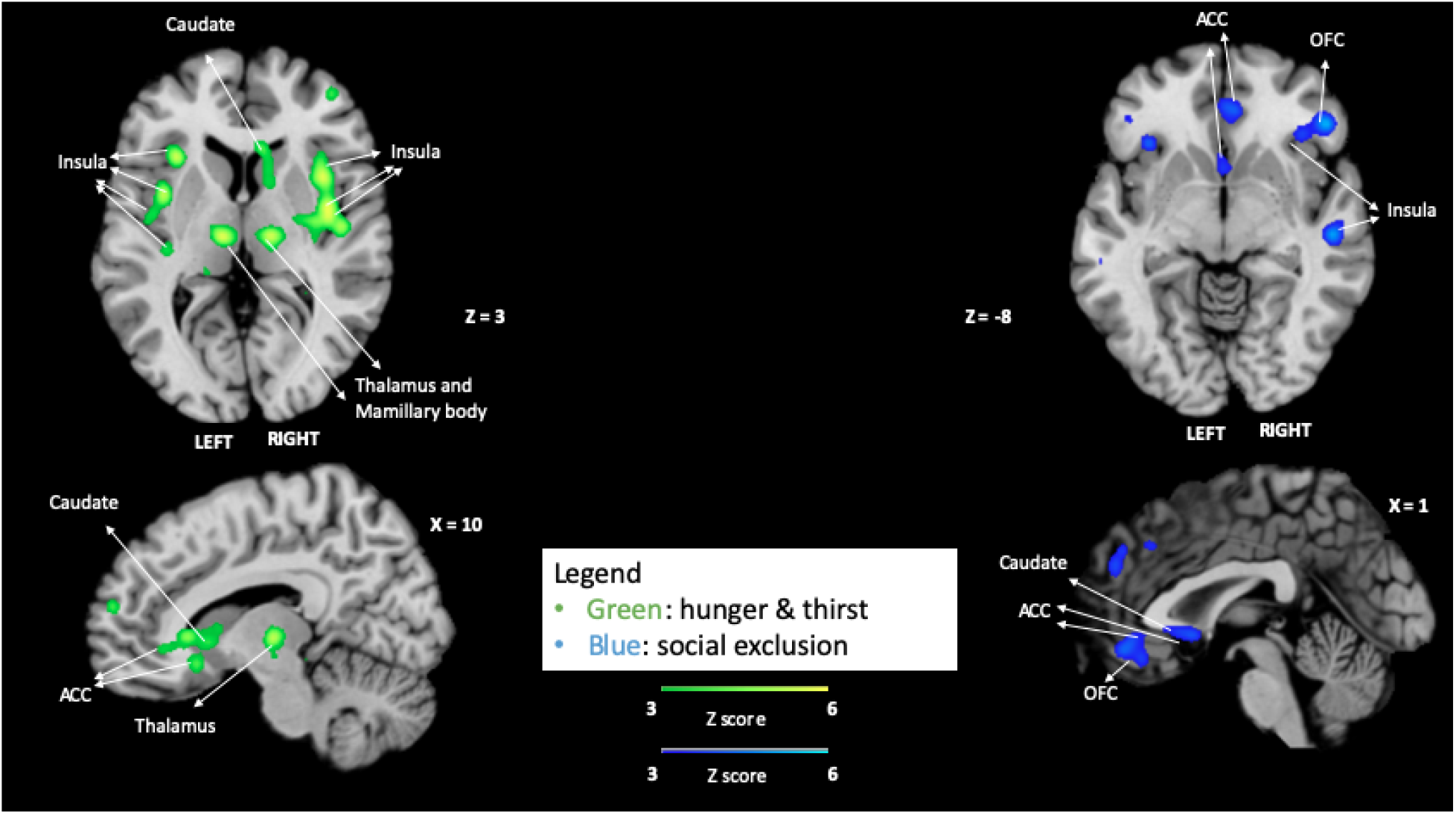
Single meta-analyses maps. Maps for activated clusters in each condition: Physiological-Need (green) and Social-Need (blue) and, showing activation patterns for each.

The single meta-analysis on Social-Need, i.e. social interaction perception during/after social exclusion, revealed consistent activation within the right anterior insula, bilateral posterior insula, bilateral ACC (B24 and B32), right inferior frontal gyrus, right OFC, left anterior transverse temporal gyrus, and bilateral caudate head (see table 6 and figure 1).

**Table 6.** Social exclusion Coordinates for peak activated clusters in the social exclusion condition. We used a cluster-level family-wise error (cFWE) correction (P < 0.05) with a cluster-forming threshold of P < 0.001(uncorrected at the voxel level) (Eklund et al., 2016; Woo et al., 2014), and 1000 permutations. cluster-level family-wise error (cFWE) correction (P < 0.05) with a cluster-forming threshold of P < 0.001(uncorrected at the voxel level) (Eklund et al., 2016; Woo et al., 2014), and 1000 permutations.

| Cluster # | x | y | z | ALE | P | Z | Label (Nearest Gray Matter within 5mm) |
| --- | --- | --- | --- | --- | --- | --- | --- |
| 1 | 40 | -16 | 18 | 0.055232868 | 3.9646563E-16 | 8.055735 | Right Cerebrum.Sub-lobar.Insula.Gray Matter.Brodmann area 13 |
| 1 | 48 | -10 | 8 | 0.021314455 | 1.0559885E-5 | 4.252716 | Right Cerebrum.Sub-lobar.Insula.Gray Matter.Brodman area 13 |
| 2 | 6 | 38 | -6 | 0.02248284 | 5.194597E-6 | 4.408918 | Right Cerebrum.Limbic Lobe.Anterior Cingulate.Gray Matter.Brodman area 24 |
| 2 | 2 | 44 | -12 | 0.021057572 | 1.2254903E-5 | 4.219271 | Right Cerebrum.Limbic Lobe.Anterior Cingulate.Gray Matter.Brodman area 32 |
| 2 | -2 | 36 | -16 | 0.019663427 | 2.7848786E-5 | 4.030337 | Left Cerebrum.Limbic Lobe.Anterior Cingulate.Gray Matter.Brodman area 32 |
| 3 | 50 | 32 | -8 | 0.027039705 | 2.7960468E-7 | 5.00482 | Right Cerebrum.Frontal Lobe.Inferior Frontal Gyrus.Gray Matter.Brodman area 45 |
| 3 | 38 | 28 | -6 | 0.020422611 | 1.7853532E-5 | 4.1336384 | Right Cerebrum.Frontal Lobe.Inferior Frontal Gyrus.Gray Matter.Brodman area 47 |
| 3 | 38 | 36 | -14 | 0.016634589 | 1.7061671E-4 | 3.5818017 | Right Cerebrum.Frontal Lobe.Inferior Frontal Gyrus.Gray Matter.Brodman area 47 |
| 4 | -36 | -16 | 20 | 0.023092672 | 3.5666646E-6 | 4.4896793 | Left Cerebrum.Sub-lobar.Insula.Gray Matter.Brodman area 13 |
| 4 | -48 | -20 | 12 | 0.022719312 | 4.485696E-6 | 4.4405947 | Left Cerebrum.Temporal Lobe.Transverse Temporal Gyrus.Gray Matter.Brodman area 41 |
| 5 | 10 | 12 | 0 | 0.02310413 | 3.5442677E-6 | 4.491021 | Right Cerebrum.Sub-lobar.Caudate.Gray Matter.Caudate Head |
| 5 | 0 | 12 | -6 | 0.019869018 | 2.4631125E-5 | 4.0591025 | Left Cerebrum.Sub-lobar.Caudate.Gray Matter.Caudate Head |
| 5 | 0 | 18 | -4 | 0.019038977 | 3.9894312E-5 | 3.945036 | Left Cerebrum.Sub-lobar.Caudate.Gray Matter.Caudate Head |

### Contrasts meta-analyses

Contrasts meta-analyses results are summarized in tables 7 and 8 and figure 2. Compared to perceiving social interaction during social exclusion, perceiving food or water during hunger or thirst elicited more consistent activation within the bilateral posterior insula, right OFC and the bilateral caudate. Compared to perception of food or water during hunger or thirst, perception of social interaction during social exclusion elicited more consistent activation within the right posterior insula, the right OFC, right inferior frontal gyrus, and left ACC (B32).

**Table 7.** [Hunger & thirst] minus [Social exclusion] Coordinates for peak activated clusters in the [Hunger & thirst] minus [Social exclusion] contrast. We used the two cFWE corrected maps with p < .01 (uncorrected at the voxel level), 10,000 permutations (see Eickhoff et al., 2011).

| Cluster # | x | y | z | ALE | P | Z | Label (Nearest Gray Matter within 5mm) |
| --- | --- | --- | --- | --- | --- | --- | --- |
| 1 | 40 | -16 | 18 | 0.055232868 | 3.9646563E-16 | 8.055735 | Right Cerebrum.Sub-lobar.Insula.Gray Matter.Brodmann area 13 |
| 1 | 48 | -10 | 8 | 0.021314455 | 1.0559885E-5 | 4.252716 | Right Cerebrum.Sub-lobar.Insula.Gray Matter.Brodmann area 13 |
| 2 | 6 | 38 | -6 | 0.02248284 | 5.194597E-6 | 4.408918 | Right Cerebrum.Limbic Lobe.Anterior Cingulate.Gray Matter.Brodmann area 24 |
| 2 | 2 | 44 | -12 | 0.021057572 | 1.2254903E-5 | 4.219271 | Right Cerebrum.Limbic Lobe.Anterior Cingulate.Gray Matter.Brodmann area 32 |
| 2 | -2 | 36 | -16 | 0.019663427 | 2.7848786E-5 | 4.030337 | Left Cerebrum.Limbic Lobe.Anterior Cingulate.Gray Matter.Brodmann area 32 |
| 3 | 50 | 32 | -8 | 0.027039705 | 2.7960468E-7 | 5.00482 | Right Cerebrum.Frontal Lobe.Inferior Frontal Gyrus.Gray Matter.Brodmann area 45 |
| 3 | 38 | 28 | -6 | 0.020422611 | 1.7853532E-5 | 4.1336384 | Right Cerebrum.Frontal Lobe.Inferior Frontal Gyrus.Gray Matter.Brodmann area 47 |
| 3 | 38 | 36 | -14 | 0.016634589 | 1.7061671E-4 | 3.5818017 | Right Cerebrum.Frontal Lobe.Inferior Frontal Gyrus.Gray Matter.Brodmann area 47 |
| 4 | -36 | -16 | 20 | 0.023092672 | 3.5666646E-6 | 4.4896793 | Left Cerebrum.Sub-lobar.Insula.Gray Matter.Brodmann area 13 |
| 4 | -48 | -20 | 12 | 0.022719312 | 4.485696E-6 | 4.4405947 | Left Cerebrum.Temporal Lobe.Transverse Temporal Gyrus.Gray Matter.Brodmann area 41 |
| 5 | 10 | 12 | 0 | 0.02310413 | 3.5442677E-6 | 4.491021 | Right Cerebrum.Sub-lobar.Caudate.Gray Matter.Caudate Head |
| 5 | 0 | 12 | -6 | 0.019869018 | 2.4631125E-5 | 4.0591025 | Left Cerebrum.Sub-lobar.Caudate.Gray Matter.Caudate Head |
| 5 | 0 | 18 | -4 | 0.019038977 | 3.9894312E-5 | 3.945036 | Left Cerebrum.Sub-lobar.Caudate.Gray Matter.Caudate Head |

**Table 8.** [Social exclusion] minus [Hunger & thirst] Coordinates for peak activated clusters in the [Social exclusion] minus [Hunger & thirst] contrast. We used the two cFWE corrected maps with p < .01 (uncorrected at the voxel level), 10,000 permutations (see Eickhoff et al., 2011).

| Cluster # | x | y | z | P | Z | Label (Nearest Gray Matter within 5mm) |
| --- | --- | --- | --- | --- | --- | --- |
| 1 | 42.4 | -16.1 | 20.8 | 2.0E-4 | 3.5400841 | Right Cerebrum.Sub-lobar.Insula.Gray Matter.Brodmann area 13 |
| 2 | 48 | 36 | -8 | 3.0E-4 | 3.4316144 | Right Cerebrum.Frontal Lobe.Middle Frontal Gyrus.Gray Matter.Brodmann area 47 |
| 2 | 44 | 36 | -8 | 4.0E-4 | 3.352795 | Right Cerebrum.Frontal Lobe.Inferior Frontal Gyrus.Gray Matter.Brodmann area 47 |
| 2 | 50 | 28 | -6 | 0.0035 | 2.6968443 | Right Cerebrum.Frontal Lobe.Inferior Frontal Gyrus.Gray Matter.* |
| 3 | -2 | 44 | -14 | 0.0026 | 2.794376 | Left Cerebrum.Limbic Lobe.Anterior Cingulate.Gray Matter.Brodmann area 32 |

**Figure 2.**
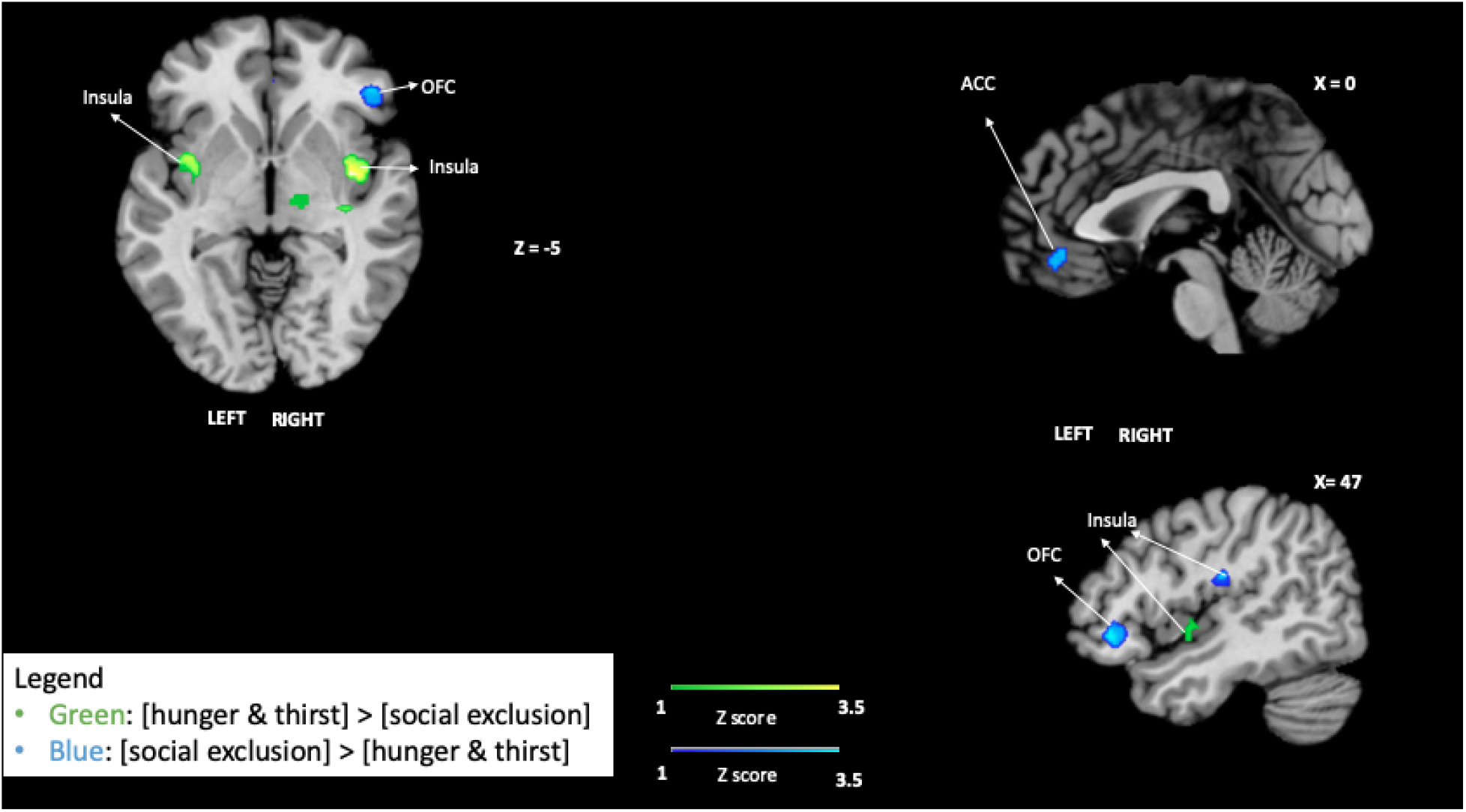
Contrasts maps. In green, clustered thresholded maps for clusters of subtraction {[Physiological-Need] minus [Social-Need]}. In blue, clustered thresholded maps for clusters of subtraction {[Social-Need] minus [Physiological-Need]}.

### Conjunction meta-analysis

The intersection between [Physiological-Need] AND [Social-Need] showed overlapping consistent activation in the right posterior insula, right caudate head, and right ACC (B24). (see table 9 and figure 3). It should be noted that the cluster size of the ACC, 8 mm^3^, is below what is usually used as minimum, i.e. 10 mm^3^.

**Table 9.** [Hunger & thirst] AND [Social exclusion] Coordinates for the intersection between s in the [Hunger & thirst] condition AND [Social exclusion] condition. the conjunction was the intersection of the two cFWE thresholded maps.

| Cluster # | x | y | z | ALE | Label (Nearest Gray Matter within 5mm) |
| --- | --- | --- | --- | --- | --- |
| 1 | 48 | -10 | 6 | 0.019197641 | Right Cerebrum.Sub-lobar.Insula.Gray Matter.Brodmann area 13 |
| 2 | 10 | 14 | -2 | 0.017616214 | Right Cerebrum.Sub-lobar.Caudate.Gray Matter.Caudate Head |
| 3 | 10 | 38 | -6 | 0.014797952 | Right Cerebrum.Limbic Lobe.Anterior Cingulate.Gray Matter.Brodmann area 24 |

**Figure 3.**
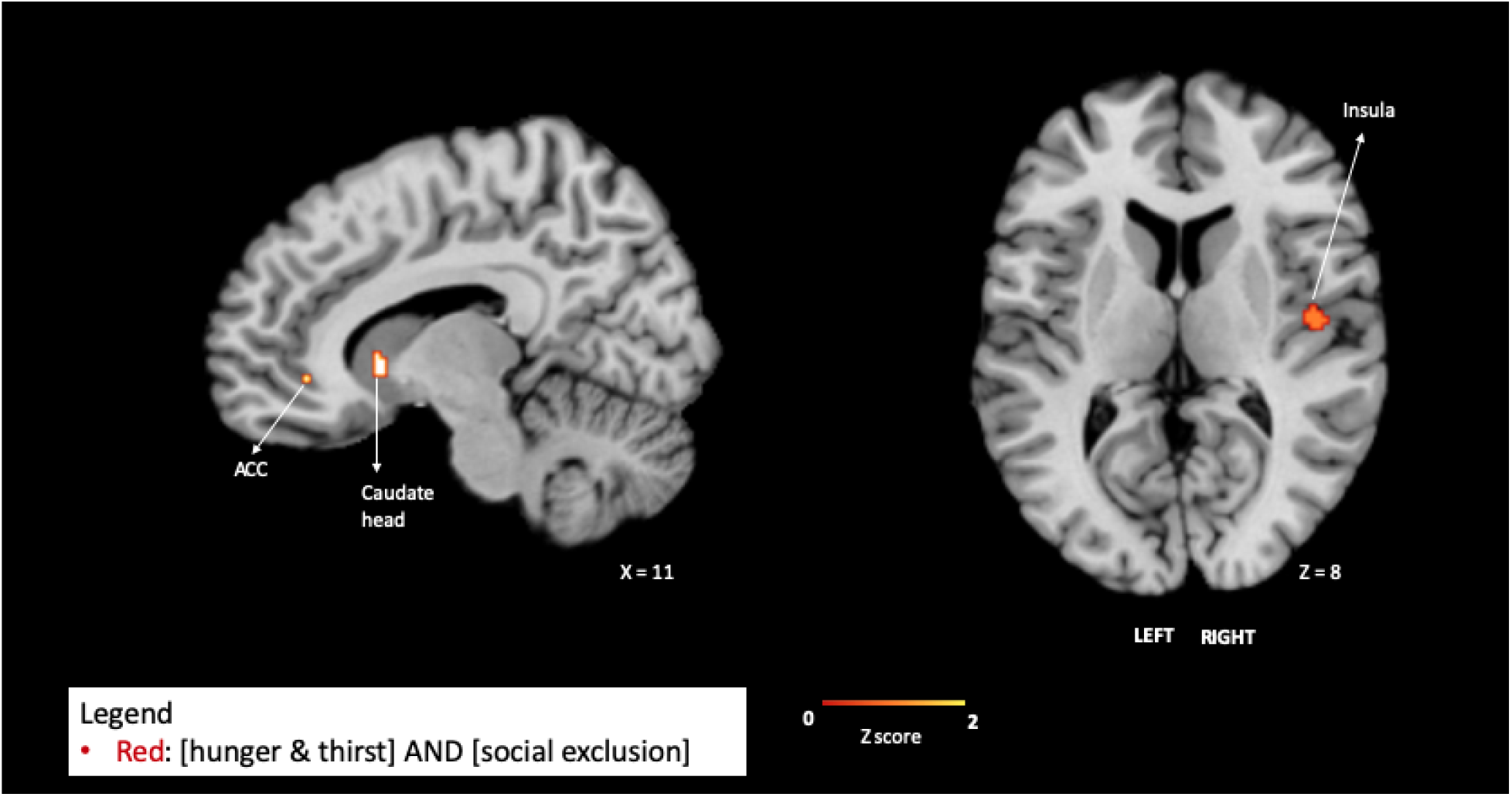
Conjunction maps. Clustered thresholded maps showing the intersection between activation patterns of [Physiological-Need] AND [Social-Need].

### Spatial correlation between the conjunction map and serotonin neurotransmitter receptors

Finally, we also looked at the topographical relationship between brain coordinates found in our conjunction analysis [Physiological-Need] AND [Social-Need] and whole-brain maps of various neurotransmitters (with a focus on serotonin receptors) available in the JuSpace. Because (low) serotonin levels in the brain have been related to sensitivity to food (van Galen et al., 2021) and social exclusion (Preller et al., 2015), our focus was on serotonin receptors. Among serotonin receptors, significant positive spatial correlation between the regions showing consistent activation for physiological and social need and neurotransmitter maps were found for 5HT4 (see figure 4). Although this was the highest correlation with serotonin receptors, it was not the highest of all receptors (see supplementary material for full results). Regarding this, positive correlations were also found for : D1 and D2 dopamine receptors, DAT dopamine transporter, VAchT acetylcholine transporter, SERT serotonin transporter, 5HT4 and 5HT1b serotonin receptors, the Mu opioid receptor, mGluR5 metabotropic glutamate receptor. We found negative correlations for 5HT2a and 5 HT1a serotonin receptors, and CB1 endocannabinoid receptors. This would indicate that dopamine, endogenous opioids, acetylcholine as well as serotonin neurotransmitters are spatially correlated with the intersection network.

**Figure 4.**
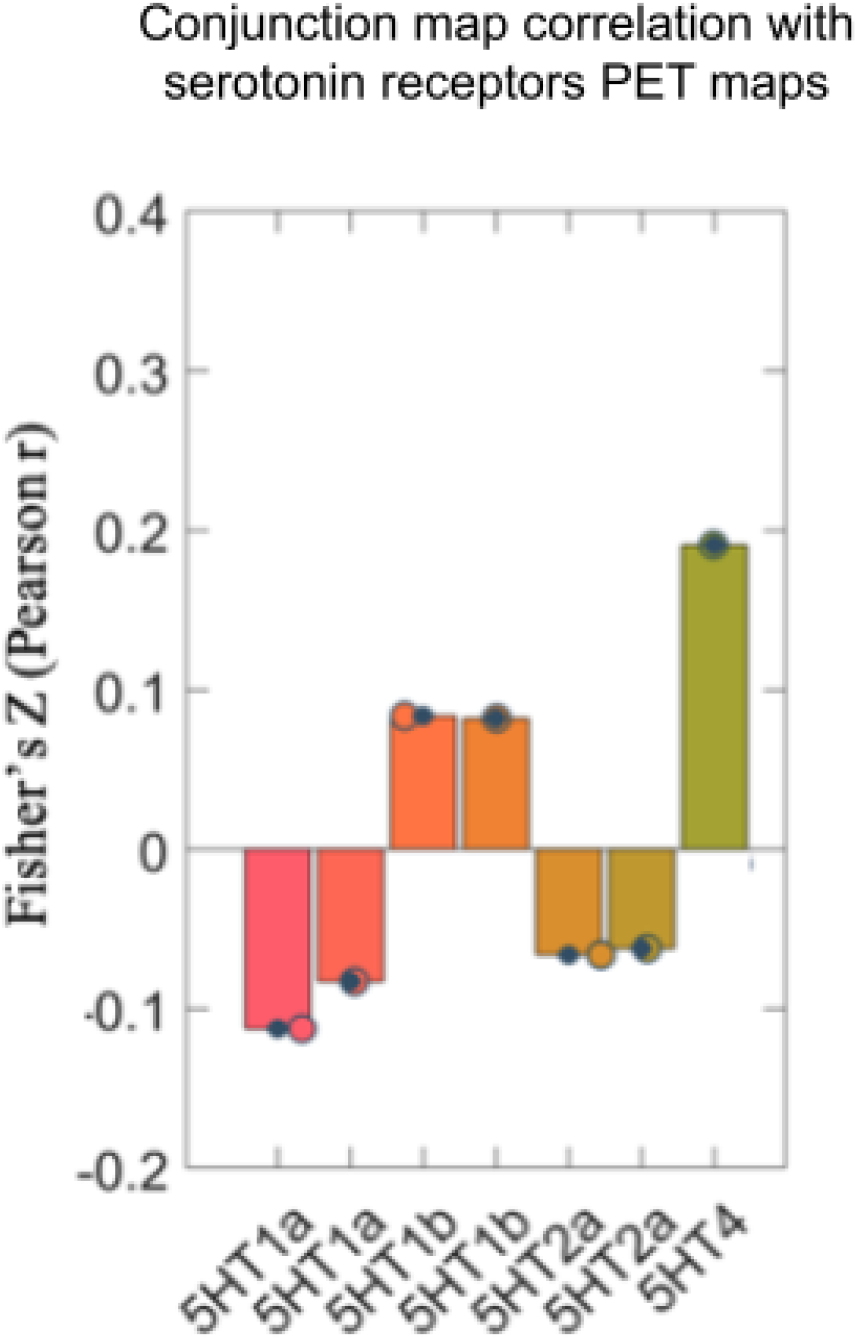
Spatial correlation between the conjunction map and serotonin receptors. Error bars showing the Fisher’s z Pearson correlation between the conjunction data on the y axis and the neurotransmitter map based on PET studies on the x axis. The colored dots represent data points and the black ones the mean of the bar which equals the to the single point.

## DISCUSSION

Our goal was to investigate possible common and specific brain activation patterns for, on one hand, the processing of physiologically (hunger and thirst) and non-physiological socially (social exclusion) needed stimuli. To achieve this objective, we used ALE neuroimaging meta-analysis, comparing consistent brain activation patterns during processing of relevant (deprived) stimuli while in physiological vs. social deprivation states. We first carried out separate single meta-analyses on physiological ‘needing’ and on social ‘needing’. For perception of food and water in hungry and thirsty states, we found consistent activity within the insula, claustrum, putamen, ACC (B24 and B25), caudate head and body, the parahippocampal gyrus, the medial frontal gyrus, amygdala, uncus, mamillary body, and hippocampus. Perception of social interaction during social exclusion revealed consistent activation within the insula, ACC (B24 and B32), inferior frontal gyrus, OFC, anterior transverse temporal gyrus, and caudate head. We then contrasted those maps to identify differences. Compared to social ‘needing’, physiological ‘needing’ more consistently elicited activity within the putamen, claustrum, anterior insula, OFC, thalamus, mammillary body and amygdala. Compared to physiological ‘needing’, social ‘needing’ more consistently activated the posterior insula, the more lateral part of OFC, inferior frontal gyrus, and ACC. We also intersected the maps of physiological ‘needing’ and social ‘needing’ to identify common brain areas between physiological and social ‘needing’. In that regard, our results suggest that both processing a physiological and social stimulus we are deprived of (i.e. ‘needing’) seems to consistently activate parts of the posterior insula, ACC and caudate. Furthermore, we used the spatial pattern of that conjunction map in order to identify a possible relation between this common network and serotonin neurotransmitter receptors. Our analysis showed that of all serotonin receptors, the 5HT4 receptor had the highest spatial correlation. In the following paragraphs, we will discuss how these results can help us understand how the brain processes relevant stimuli in a deprived state.

Our single meta-analyses results are in line with the literature. Specifically, previous studies reported activation of insula and ACC for perception of food when hungry (van der Lan et al., 2011; Goldstone, et. al., 2009), water when thirsty (De Araujo et al., 2003), and social interaction when excluded (Mwilambwe & Spreng, 2021). Similar to these, we found consistent activation within the ACC and insula during perception of the physiologically and socially needed stimuli in our single meta-analyses. In that sense, our results show that there is indeed an overlap between processing of physiologically needed stimuli and socially needed stimuli. The contrast [Physiological-Need] minus [Social-Need] revealed activation within the posterior insula, dorsal ACC, pregenual ACC, OFC, and caudate head, whereas the contrast [Social exclusion] minus [Hunger & thirst] did not include the caudate, but also revealed activation within the posterior insula, a more lateral part of the OFC and pregenual ACC. Our single and contrast meta-analyses suggest that neural populations treating physiologically and/or socially needed stimuli might be spatially closer (or even be the same) in the posterior insula and pregenual ACC. However, for other regions, (such as the OFC) that seem to be activated by both types of need when looking at a macro level, the activations are in different sub-regions when taking a closer look.

States such as hunger and thirst are referred to as homeostatic emotions (Craig, 2003). Despite how they are generated, homeostatic emotions have two important components : the aversive affect, and the affective motivation to terminate that affect (Craig, 2003). More specifically, thalamocortical projections provide both information about (1) the physiological condition of the body in interoceptive cortex at the dorsal and posterior part of the insula; as well as (2) activation of limbic motor cortex, i.e., the ACC (Craig, 2003). These respectively generate the affective perception and motivation components (Craig, 2003). This is in line with our single meta-analysis results on hunger/food and thirst/water showing consistent activation within the posterior insula and the ACC.

Social exclusion has been said to activate the same dorsal region within the ACC as physical pain (Eisenberg, 2012). However, a recent meta-analysis of social exclusion with the cyberball task found a more ventral part of the ACC (Milambwe & Spreng, 2021). This latter result is similar to our single meta-analysis findings. So, it’s possible that social exclusion and physical pain have slightly different brain activity patterns. Also, though social exclusion has been linked to activation of the anterior insula rather than the posterior insula (Eisenberg, 2012), our single meta-analysis on social exclusion found activity in both the posterior insula and in the anterior insula. This is in line with Milambwe & Spreng (2021) recent meta-analysis on social exclusion. However, Vijayakumar and colleagues (2017) did not find the insula in their social exclusion meta-analysis. The difference between our findings and Vijayakumar and colleagues (2017) findings might be related to technical issues or inclusion criteria. Moreover, our study went further than the preceding meta-analyses by investigating the brain activation pattern between perception of social interaction while excluded and that of food/water when hungry/thirsty, enabling us to further assess the need aspect of social excursion in terms of brain patterns.

Some unanswered questions regarding need states might be hinted by looking at the common brain activation pattern map between psychologically and socially needed stimuli. Regarding this, our study shows that consistent brain activity patterns during perception of social interaction while being excluded partly intersects with that of perception of physiologically needed stimuli. This intersection was found within the posterior insula, ventral ACC and caudate which are related to aversive affects, affective motivation and goal directed behavior, respectively (Craig, 2003; Loonen & Ivanova., 2018; Balleine and O’Doherty, 2010). These three regions might be working in the following way: (1) An aversive affect perception of ‘needing’ is likely processed within the dorsal posterior insula (Craig, 2003), which integrates aversive emotional states and other homeostatic bodily functions (Gerlach et al., 2019) and likely signals a difference between actual and desired interoceptive state (Livneh et al., 2020; Barrett & Simmon, 2015). (2) The ACC contributes in facilitating directional motivation (Craig, 2003) possibly signaling the fulfillment of ‘needing’ as a requirement and distributing that signal to other brain regions (Weston, 2012) including the caudate (Peak et al., 2019). (3) Activation of the latter might indicate that perceiving needed stimuli leads to goal directed behavior and action choice (Balleine and O’Doherty, 2010; Knutson & Cooper, 2005; Hollon et al., 2014; ito and Doya, 2015; Schwabe & Wolf, 2010), based on the current need (van den Bercken & Cools, 1982). Thus, our method and idea of looking at the common brain areas between psychologically and socially needed stimuli allowed us to further elucidate this mechanism. Following that, our findings suggest that perception of both physiologically and socially needed stimuli embed the affective perception, processed within the posterior insula, and the affective motivation toward the stimuli/event that terminates that need state, processed within the ventral ACC, which directs action choice within the caudate (Peak et al., 2019). Though this was initially suggested for homeostatic/physiological needs and their relevant stimuli (see Craig, 2003), our results lead us to suggest that this might be also true for social need states and social interaction.

As discussed in the previous paragraph, a possible link between social exclusion and hunger/thirst could be that they are both related to some displeasure/aversive state. This might also explain the inconsistency in the relationship between social exclusion and pain (Eisenberg, 2012; Milambwe & Spreng, 2021): social exclusion might be related, not to pain, but to displeasure. The latter is the inverse of pleasure (Cabanac, 2002) and as such includes, but is not limited to, pain. Indeed, displeasure can also be related to hunger, shortness of breath, disgust, depression, anxiety, fear, etc. (Becker et al., 2019; Berridge & Kringelbach, 2015) and is said to be processed within the pregenual ACC (and mesolimbic and amygdalar circuitry) (Becker et al., 2019), a region that we identified as being commonly recruited during both physiological and social needs.

It is possible that physiological needs and social exclusion share the negative feeling aspect of needing, and not as much the motivational aspect. Whereas, physiological needs and social isolation might share such a motivational aspect. Indeed, a recent study by Tomova and colleagues (2020) on the common brain areas between perception of food during hunger and social interaction after social isolation only found common activation within the substantia nigra (SN) and ventral tegmental area (VTA). These are dopaminergic regions, and the VTA mesolimbic dopamine has been linked to reward prediction, reward learning and motivation (Schultz et al., 1997; Schultz, 2015; Montague et al., 1996; Schultz, 1998; Rice et al., 2010; Hamid et al., 2016). However, in our conjunction analysis, which was about social exclusion rather than isolation, we did not find the VTA/SN nor the ventral striatum which receives VTA dopamine for action invigoration and reward seeking (Li and Daw, 2011; Berridge and Aldrige, 2009; Lex and Hauber, 2008; Hamid et al., 2016; Zhang et al. 2009; Balleine and Killcross, 2006). This is an indication that the difference between our findings and Tomova et al. (2020)’s is due to the difference between social exclusion and social isolation. This has been confirmed by other studies on social isolation (see Inagaki et al., 2016). However, it is important to note that not all studies on social isolation have found activity within the mesolimbic dopaminergic VTA or ventral striatum (see Cacioppo et al., 2009; D’agostino et al., 2019).

Our aim was to go beyond common brain activity patterns between psychologically and non-physiological socially needed stimuli, by running exploratory analysis to look at how these common brain patterns–posterior insula, caudate and ACC–might be related to serotonergic receptors distribution in the brain. Our results show that at the conjunction regions between physiological and social ‘needing’, the serotonin receptor with highest density is the 5HT4 receptor. Our finding is in line with the suggestion that the 5HT4 receptor is a component of a feedback loop from the medial PFC to the dorsal raphe nuclei (DRN) (Rebholz, et al., 2018), specifically the prelimbic and infralimbic subregions (Peyron et al 1998, Lucas et al, 2005), which correspond to Brodmann areas 32 and 25 respectively (Price, 2007); which are part of the ACC (Weston, 2012). The ACC activity is related to both an affective motivation and an update of internal models, i.e. a feedback loop (Craig, 2003; Kolling et al., 2016; Petzschner et al., 2021). Some information from this feedback loop is said to be sent from the ACC to the dorsal raphe nucleus (DRN) (Rebholz et al., 2018; Lucas et al., 2005) which is the largest serotonergic structure in the brain (Liu et al., 2020). Our finding related to the spatial correlation between 5HT4 and the common brain map, leads us to suggest that this feedback loop can allow need states or needed stimuli to influence activity in the DRN through the ACC. That influence can be either inhibitory or excitatory via modulation of GABAergic (DRN inhibition) neurons and CB1 receptors (DRN excitation) in the DRN by the ACC (Lucas et al., 2005; Castello, et al 2018; Geddes et al.2016). Indeed CB1 has been found to be implicated in pleasure for both food (Kirkham, 2009) and social play (Achterberg et al., 2016). This might give some reason why needed stimuli are pleasurable. However we did not find much correlation between the common pattern map and CB1; suggesting that that CB1 actions could be further downstream, closer to the DRN rather than closer to the common brain map found in this study: posterior insula, pregenual ACC and caudate. Furthermore, though our findings did not show activity in the mesolimbic dopaminergic motivational areas, e.g. SN/VTA and ventral striatal activation; our finding, that the intersection network is correlated to 5HT4 distribution, could partly explain how needing, physiological or social needs can lead to activity within those areas. Indeed, the VTA\SN is modulated by DRN (Gervais & Rouillard, 2000), whose activity, as our study and literature suggest, is regulated by ACC via the 5HT4 receptor. In that sense: via 5HT4, pregenual ACC regulates the serotonergic DRN, which in turn can influence VTA/SN dopaminergic activity. In summary, our study suggests that the perception of a stimulus that would alleviate a negative physiological or social state (need) could be linked to brain regions that influence the activity of serotonin neurons.

## LIMITS

The main limit of our study is that for physiological needs we only included hunger and thirst, while for social needs we only included social exclusion and not isolation. Nevertheless, it has been argued that both hunger and thirst implicate a similar network, that includes the ACC and insula (Mckinley et al., 2019), and the mechanism of that network is similar to other needs related to temperature, itch, visceral distension, muscle ache, ‘air hunger’, etc. (Crag, 2003). In the same way, social exclusion is different from social isolation: social isolation is more related to meaningful social contacts whereas exclusion is more related to being outcast of, or not able to participate in, some society (Huisman & van Tilburg, 2021)). Moreover, the contrasts of social exclusion included here are short term non physiological social needs, whereas social isolation is more long term, in terms of hours, and can have physiological impacts (See Tomova et al., 2020; Cacioppo et al., 2002). However they still have in common the fact that they refer to “lack of ties”, either with with society (exclusion) or with other significant persons (isolation) (Huisman & van Tilburg, 2021). Also, although there has been a debate between whether social exclusion actually causes need related distress (Gerber & Wheeler, 2009; Blackhart et al., 2009), Bernstein and Claypool (2012) found that Cyberball exclusion paradigm, used in this paper, may induce need-like emotional distress (e.g., reduced mood, and lowered self-esteem and other needs). Hence, the use of cyberball in this study is more coherent with a need state (Bernstein & Claypool, 2012). Future studies could assess a larger range of physiologically and socially needed stimuli, including those not included in the present study. Furthermore, one should not forget the limitations of reverse inference (Poldrack, 2006; 2011) in interpreting our results.

## CONCLUSION

Our goal was to study the common and specific brain activations during physiological (hunger and thirst) and non physiological social (social exclusion) ‘needing’ as well as their relationship to the serotonergic system. Our results suggest that regions within the mid-posterior insula, the ACC and the caudate are regions that commonly support processing/perception of both physiologically (hunger and thirst) and socially (exclusion) needed stimuli. So our result lead us to propose that ‘needing’ whether physiologically or socially is related to (1) am affective perception or response towards the needed stimulus that signal difference between actual and desired state, and which is processed within in the mid-posterior insula; (2) an affective and directional motivation that requires the termination of the need state, processed within the ACC. (3) This requirement to terminate the need state facilitates goal directed behavior within the caudate. Furthermore, The network of regions at the intersection seem to be related to the distribution of receptors, and among the serotonergic receptors, the 5HT4 seem to have one the highest spatial correlation with that network. We hypothesize that, in need state and/or while processing the needed stimuli, this intersection network, through 5HT4, modulates DRN serotonin activity which signals how beneficial, or not, the current state is. In that sense, in the brain, physiological and social deprivation could lead to low serotonin levels, whereas the onset or presence of physiologically and socially needed stimuli could be related to high serotonin levels.

## Supporting information

PRISMA

## ACKNOWLEDGMENTS

The research was supported in part by NSERC Discovery Grant #RGPIN-2018-05698 and UdeM institutional funds.

## AUTHOR CONTRIBUTION

**Juvénal Bosulu**: Designed the study, performed the database search, performed data analysis, interpretation, and wrote the manuscript. **Sébastien Hétu**: Revised the manuscript and provided critical feedbacks. **Yi Luo**: Revised the manuscript and provided critical feedbacks. All authors contributed to and approved the final manuscript version.

## DATA AVAILABILITY STATEMENT

All Data are available upon request.

