## Supplementary material for "Common brain areas for processing physiologically and socially ‘needed’ stimuli": PRISMA

**PRISMA Hunger**

Studies included in quantitative synthesis (meta-analysis)
(n = 16)

Studies included in qualitative synthesis
(n = 16)

Full-text articles excluded, with reasons
(n = 25)

Full-text articles assessed for eligibility
(n = 41)

Records excluded
(n = 116)

Records screened
(n = 157)

Records after duplicates removed
(n =	157)

Additional records identified through other sources
(n = 7)

### Identification

### Eligibility

### Included

### Screening

Records identified through database searching
(n = 150)

**PRISMA for thirst**

Studies included in quantitative synthesis (meta-analysis)
(n = 4)

Studies included in qualitative synthesis
(n = 4)

Full-text articles excluded, with reasons
(n = 7)

Full-text articles assessed for eligibility
(n = 4)

Records excluded
(n = 11)

Records screened
(n = 11)

Records after duplicates removed
(n =	22)

Additional records identified through other sources
(n = 3)

### Identification

### Eligibility

### Included

### Screening

Records identified through database searching
(n = 19)

**PRISMA for Social exclusion/isolation**

Studies included in quantitative synthesis (meta-analysis)
(n = 26)

Studies included in qualitative synthesis
(n = 26)

Full-text articles excluded, with reasons
(n = 28)

Full-text articles assessed for eligibility
(n = 26)

Records excluded
(n = 79)

Records screened
(n = 54)

Records after duplicates removed
(n =	133)

Additional records identified through other sources
(n = 4)

### Identification

### Eligibility

### Included

### Screening

Records identified through database searching
(n = 129)
